# Green synthesis of ZnO nanoparticles using *Cocos nucifera* leaf extract: Characterization, antimicrobial, antioxidant, and photocatalytic activity

**DOI:** 10.1101/2022.10.27.514023

**Authors:** Farjana Rahman, Md Abdul Majed Patwary, Md. Abu Bakar Siddique, Muhammad Shahriar Bashar, Md. Aminul Haque, Beauty Akter, Rimi Rashid, Md. Anamul Haque, A. K. M. Royhan Uddin

## Abstract

Zinc oxide nanoparticles (ZnO NPs) have been successfully prepared using *Cocos nucifera* leaf extract and investigated with their antimicrobial, antioxidant, and photocatalytic activity. The structural, compositional, and morphological properties of the NPs were recorded and studied systematically to confirm the synthesis. The aqueous suspension of NPs showed a UV-Vis. absorption maxima of 370 nm indicating primarily its formation. The XRD analysis identified the NPs with hexagonal wurtzite structure with an average particle size of 16.6 nm. The FTIR analysis identified some biomolecules and functional groups in the leaf extract as responsible for the encapsulation and stabilization of ZnO NPs. The EDX analysis showed the desired elemental compositions in the material. A flower-shaped morphology of ZnO NPs was observed by SEM with a grain size of around 15 nm. The optical properties of the NPs were studied by UV-Vis spectroscopy and the band gap was calculated as 3.37 eV. The prepared ZnO NPs have demonstrated antimicrobial activity against *T. harzianum and S. aureus* with a ZOI (zone of inhibition) of 14 and 10 mm, respectively. The photocatalytic behavior of ZnO NPs showed absorbance degradation at around 640 nm and discolored methylene blue dye after one hour with a degradation maximum of 84.29 %. Thus, the prepared ZnO NPs could be used potentially in antibiotic development, pharmaceutical industries, and as photocatalysts.

## 1. Introduction

Nanoscience and nanotechnology are the most emerging fields in recent times and moving forward sharply along with physics, chemistry, biology, molecular engineering, and so on. Nanomaterials are in versatile use in pharmaceutical, cosmetic, textile, and even electrical and electronics industries. Nanomaterials are products processed through nanotechnologies that contain nanoparticles (NPs) on a scale ranging from 1 to 100 nm. The NPs of metal and metal oxides are usually used in industries as requirements. Several types of metal and metal oxide NPs such as aluminum, nickel, silver, copper, copper oxide, iron, iron oxide, cerium dioxide, titanium dioxide, and zinc oxide are commonly known [1, 2]. The NPs can be prepared by several methods like physical, chemical, and biological, but physical and chemical methods are associated with high energy demand, sometimes generating poisoned and parlous chemicals, which may lead to consanguineous menaces [3, 4]. To minimize these problems a safe, cost-effective, and less hazardous synthesis procedure is already developed by modern scientists which are called the biological or green method by using plant extract with a little concentration of the chemicals. Among all metal oxides, ZnO NPs have drawn more attention for their safe and inexpensive production and preparation process [5, 6]. ZnO has been enrolled as one of the safest metal oxides by the US FDA (Food and Drug Administration) [7]. There are a lot of applications of ZnO in engineering, biological, and medicinal fields. ZnO NPs have several engineering applications such as in solar cells [8–10], gas sensors [11], chemical sensors [12], biosensors [13], and photodetectors [14], whereas, in biological and medicinal applications, ZnO NPs have cytotoxic activity [15], antimicrobial and fungicidal activities [16], anti-inflammatory activity, capability to quicken wound healing, antidiabetic [17, 18], and chemiluminescent properties [19, 20].

Current studies have supported the synthesis of ZnO NPs in several nanosized from various plant parts like leaf, flower, seed, fruit, root, rhizome, stem, bark, shell, and peel extracts. For example, some researchers utilized the leaf extracts of *Pandanus odorifer* [21], *Eucalyptus globulus* [22], *Aloe barbadensis* [23], *Sechium edule* [24], *Saponaria officinalis*[25], *Annona squamosal*[26], *Artocarpus heterophyllus*[27], *Mangifera indica* [28], *Laurus nobilis* [29], flower extracts of *Trifolium pretense* [30], *Anchusa italic* [31], *Punica granatum* [32], seed extracts of *Cuminum cyminum* [33] *Pongamia pinnata* [34], fruit extracts of *Emblica Officinalis* [35] *Borassus flabellifer* [36] *Artocarpus gomezianus* [37], root extracts of *Rubus fairholmianus* [38] *Withania somnifera* [39], rhizome extracts of *Zingiber officinale* [40] *Bergenia ciliate* [41], stem extracts of *Phyllanthus embilica* [42], bark extracts of *Cinnamomum verum* [43], *Albizia lebbeck* [44] as well as peel extracts of *Punica granatum* [45], *Musa sapientum* [46], and so on.

Previously, we have reported the green synthesis of Ag NPs for enhanced antibacterial activity using *Cocos nucifera* leaf extract [47]. As a continuation of this work, this study illustrates the green synthesis of ZnO NPs using *Cocos nucifera* leaf extract with profound antimicrobial, antioxidant, and photocatalytic activity. *Cocos nucifera* is a perennial tree that grows in tropical seashore areas, the plant is about ~100 ft long, and its leaf is about ~13 ft [48]. This plant grows best in high rainfall areas and soils with pH (5.5-7) [49] and has various medicinal uses and properties, for example, antidiarrheal, antirheumatic, aphrodisiac, cytotoxic, diuretic, emetic, emollient, hypotensive, kidney treatment, poultice, and vermicide properties [50]. *Cocos nucifera* is a fascinating plant with diverse usages ranging from domestic to therapeutic. In the endosperm (coconut meat), endocarp (coconut hard shell), and leaf extract, the existence of phytochemicals such as tannin, saponin, alkaloid, phenol, flavonoid, and volatile oil was determined. Only in the case of the leaf extract of the plant, alkaloid, tannin, saponin, and flavonoid were identified [51]. Functional groups such as alkaloids are the phytochemicals, that act as capping and reducing agents to prepare ZnO NPs [52, 53] using Zn-salt e.g. Zn(NO_3_)_2_.6H_2_O.

Several research works have already been done on *Cocos nucifera*. Roopan et al. [54] studied phytoconstituents, biotechnological applications, and nutritive aspects of Coconut (*Cocos nucifera*). Satheshkumar et al. [55] used curry leaves extracted with coconut water to synthesize ZnO-NPs and observed photocatalytic dye degradation and antibacterial activity. Priyatharesini et al. [56] used *Cocos nucifera* male flower extract to synthesize ZnO NPs and analyzed their antimicrobial activity. Krupa [57] used the endosperm of *Cocos nucifera* (coconut water) to synthesize ZnO NPs and studied tetraethoxysilane (TEOS) sol-gel coatings for combating microfouling. But, still, there is no report on the green synthesis of ZnO NPs using *Cocos nucifera* leaf extract. In this work, we have developed a *Cocos nucifera* leaf extract-mediated facile, cost-effective, and green approach for the preparation of ZnO NPs. The structural, morphological, and optical properties of the NPs are well explored. Finally, the antimicrobial, antioxidant, and photocatalytic activities of the prepared ZnO NPs are studied successfully with potent outcomes.

## 2. Materials and Methods

### 2.1. Chemicals and reagents

*Cocos nucifera* fresh leaves were collected from the local area of Cumilla, Bangladesh. Reagent grade (purity ≥ 98 %) Zn(NO_3_)_2_.6H_2_O and NaOH pellets were purchased from Fluka Analytical, Sigma-Aldrich, Germany. Mueller–Hinton Agar and potato dextrose agar were purchased from HiMedia Laboratories Pvt. Ltd., Mumbai, India. Methylene blue dye, methanol, 2, 2-diphenyl-1-picrylhydrazyl (DPPH), and ascorbic acid were purchased from Merck, Germany. All reagents and chemicals were utilized as received with no further purifications.

### 2.2. Preparation of leaf extract

The fresh leaves of *Cocos nucifera* were washed several times by using deionized water to dispel dirt particles. After washing the leaves were left to sun dry and grind to fine powder by a mortar. The fine powder leaves (about 5 g) were taken in a 250 mL beaker, mixed with 50 mL of deionized water, and heated at 80 °C for 20 minutes. Then, the mixture was filtered in another beaker with Whatman no.1 filter paper and the extract was formed in this stage according to literature [47, 58–60]. The extract was then cooled down and stored in the refrigerator (4 °C) for utilization in the synthesis of ZnO NPs, as demonstrated in Fig. 1.

**Fig. 1.**
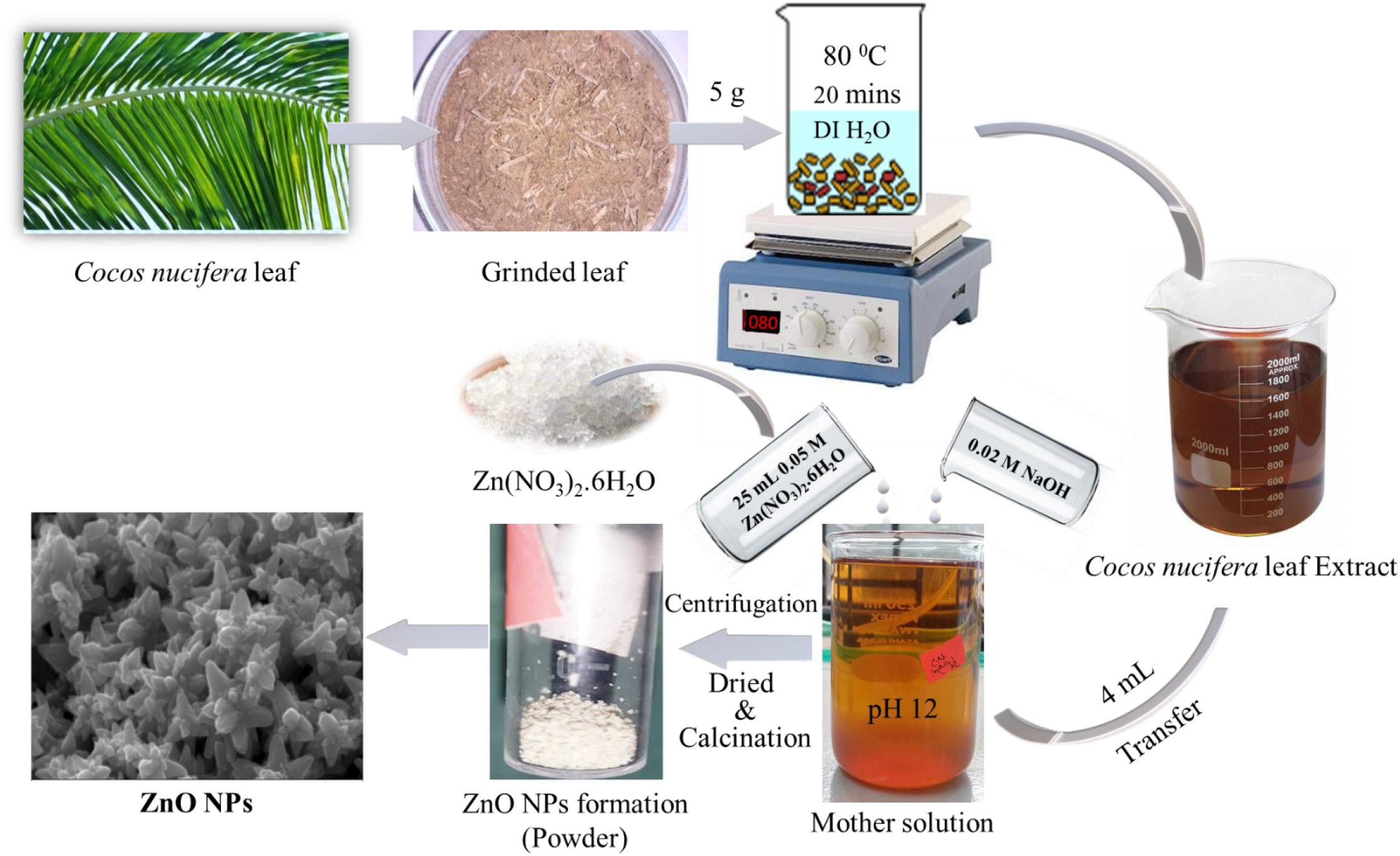
Preparation of ZnO NPs using *Cocos nucifera* leaf extract

### 2.3. Preparation of ZnO Nps

For the preparation of ZnO NPs, 25 mL of 0.05 M aqueous solution of Zn(NO_3_)_2_.6H_2_O was mixed with 4 mL of the prepared aqueous leaf extract of *Cocos nucifera* in a 250 mL beaker as demonstrated in Fig. 1 [47, 58–60]. Then, the pH of the mixture was adjusted to 12 by the drop-wise addition of 0.02 M aqueous NaOH solution. The total solution, known as the mother solution, was stirred for almost three hours with a magnetic stirrer at ambient temperature and then centrifuged by a high-speed Benchtop centrifuge machine (model no. H3-18K, Kecheng, China) at 8000 rpm for 20 minutes. This results a solid product with light-orange color which was then dried overnight in an oven (model no. LDO-060E, Labtech, Korea) at 60 °C. After that, the solid product was collected. Finally, the light-orange colored product turns into white powder when it was calcined in a muffle furnace (model no. FTMF-703, SCI FINETECH, Korea) at 550 °C for 30 minutes. The white powder was collected in a small sample vial after cooling and stored in a desiccator for further use.

### 2.4. Characterization techniques of the prepared ZnO NPs

To characterize the prepared ZnO NPs, we have utilized diverse analytical tools such as Ultraviolet-Visible (UV-Vis) spectroscopy, X-Ray Diffraction (XRD) analysis, Fourier Transform Infrared (FTIR) spectroscopy, Energy Dispersive X-Ray (EDX) spectroscopy, and Scanning Electron Microscopy (SEM). UV-Vis spectrophotometer (UV prove 1800, Shimadzu, Japan) was used primarily to confirm the formation of ZnO NPs, and the spectrum was recorded in the range of 300-500 nm by utilizing deionized water as a reference. The XRD was used to identify the phase and information of unit cell dimensions of the prepared materials [61]. The XRD of the powdered ZnO NPs was conducted by Explorer GNR, using monochromatic Cu Ka radiation (1.5419 Å) operated at a voltage of 40 kV and current of 30 mA with 2θ angle (30° – 80°) pattern and scan seed of 2°/minutes. All possible diffractions are determined by scanning the sample from 2θ angles due to the casual orientations of the powder sample. These diffraction peaks are converted to d-spacings which allow the identification of materials which is specific for each material [62]. The estimation of crystalline size (D) of the prepared ZnO NPs was calculated by Debye–Scherrer formula [63] shown below.

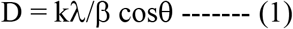

Where k (value 0.9) is the shape factor (dimensionless), λ is the X-ray wavelength of 1.5419 Å, β is the Full Width at Half Maximum (FWHM) in radian, and θ is the Bragg’s angle in radian.

The FTIR spectrophotometer (IRAffinity-1S, Shimadzu, Japan) was employed to identify the presence of the characteristics of functional groups coming from the conjugation between nanomaterial and the adsorbed biomolecules [64]. The FTIR spectrum of the prepared powdered sample was recorded in a wide range of wavenumbers (400-4000 cm^−1^) with 20 no. of scans and 2 cm^−1^ resolution which utilized the Happ-Genzel apodization function and KBr pellet method. The surface morphology and elemental composition of the prepared material were studied by SEM equipped with EDX (Model: EVO18, Carl Zeiss Microscopy, USA). A high-quality surface image of the sample was obtained by scanning the material surface with a focused beam of electrons from an electron gun applying an acceleration voltage of 15 kV.

### 2.5. Antimicrobial screening of the prepared ZnO NPs

Antimicrobial screening is an important method of analysis of the inhibitory effects of compounds against microorganisms [65]. There are a few laboratory methods available to evaluate the antimicrobial activity of a compound. The agar dilution or disc diffusion method is the most common of all methods [66]. Antimicrobial screening of ZnO NPs (50 μL dose used & concentration was 150 μg disc^−1^) was assessed against various bacterial and fungal strains. Three gram (+ve) bacterial strains such as *Staphylococcus aureus* (cars-2), *Bacillus megaterium* (*BTCC-18*), and *Bacillus cereus* (carsgp-1) as well as two fungal strains, for instance, *Aspergillus niger (carsm-3)* and *Trichoderma harzianum (carsm-2)*) were used in agar well diffusion method alike to our previous report [67]. Mueller–Hinton Agar (HiMedia, India) was used to form agar medium to culture bacteria and potato dextrose agar medium (HiMedia, India) was used to culture fungal strains. *Ceftriaxone* (10 μL) for bacterial strains and *amphotericin-B* (10 μL dose and 50 μg disc^−1^) for fungal strains were used as standards [67]. After placing the sample in a culture medium, the discs were incubated for 24 h at 37 °C for bacteria and 48 h at 26 °C for fungi in the incubator. Measuring the zone of inhibition (ZOI), the antimicrobial activity was determined.

### 2.6. Photocatalytic behavior of ZnO NPs

The ZnO NPs can act as photocatalysts because they exhibit photocatalytic activity under irradiation of sunlight [5, 24]. For this experiment, we used a 50 mg/L aqueous solution of methylene blue (MB) dye with a 5 mg/20 mL catalytic load of ZnO Nps. After mixing both solutions (dye and catalyst), the blue color of the dye solution turns into a colorless solution within 1 hour. The UV absorption was taken after 0, 10, 15, 30, and 60 minutes. The percentage of degradation was calculated by the following equation.

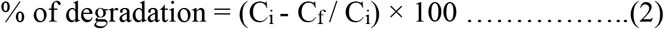

Where C_i_ and C_f_ are the initial and final concentrations of dye that degrade with time.

### 2.7. DPPH radical scavenging activity assay

2, 2-diphenyl-1-picrylhydrazyl (DPPH) radical scavenging assay was done to determine the ability of the prepared ZnO NPs to scavenge free radicals. The ability of NPs to inhibit oxidation was tested by decolorizing a methanol solution of DPPH. In methanol solution, DPPH creates a violet/purple color, which fades to shades of yellow in the presence of antioxidants. A 0.1 mM DPPH in methanol solution was prepared and 2.4 mL of it was combined with 1.6 mL of extract in methanol at varying concentrations (6.25-1200 μg/mL). The reaction mixture was vortexed completely and kept at room temperature for 30 minutes in the dark. At 517 nm, the absorbance of the mixture was determined spectrophotometrically. Ascorbic acid was utilized as a standard. The following equation was used to compute the percentage of DPPH radical scavenging activity.

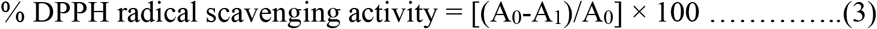

where, the absorbance of the control is A_0_, and the absorbance of the sample is A_1_.

The percent of inhibition was then plotted against concentration, and the IC_50_ was derived from the graph. At each concentration, the experiment was performed three times [68].

## 3. Results and discussions

### 3.1. Mechanism of ZnO NPs formation

There are several proposed mechanisms for the formation of ZnO NPs in the green synthesis approach [21–46, 69, 70]. In the present work, we have used the aqueous extract of *Cocos nucifera* leaf as the stabilizing as well as the natural reducing agent for ZnO NPs preparation. The phytochemical screening of this leaf demonstrated the existence of various phytochemicals such as alkaloids, resins, steroids, glycosides, terpenoids, flavonoids, polyphenols, and aromatic hydrocarbons [52, 53]. The presence of these phytochemicals in *Cocos nucifera* leaf plays an important role in the NPs preparation acting as a reducing and capping agent. **Fig. 2** showed the probable reaction mechanism for the formation of ZnO NPs in which aromatic hydroxyl groups present in the phytochemicals and polyphenols are attached with the Zn^2+^ ions from Zn(NO_3_)_2_.6H_2_O to form a stable complex system. This complex system release ZnO NPs after centrifugation and calcination [71–73].

**Fig. 2.**
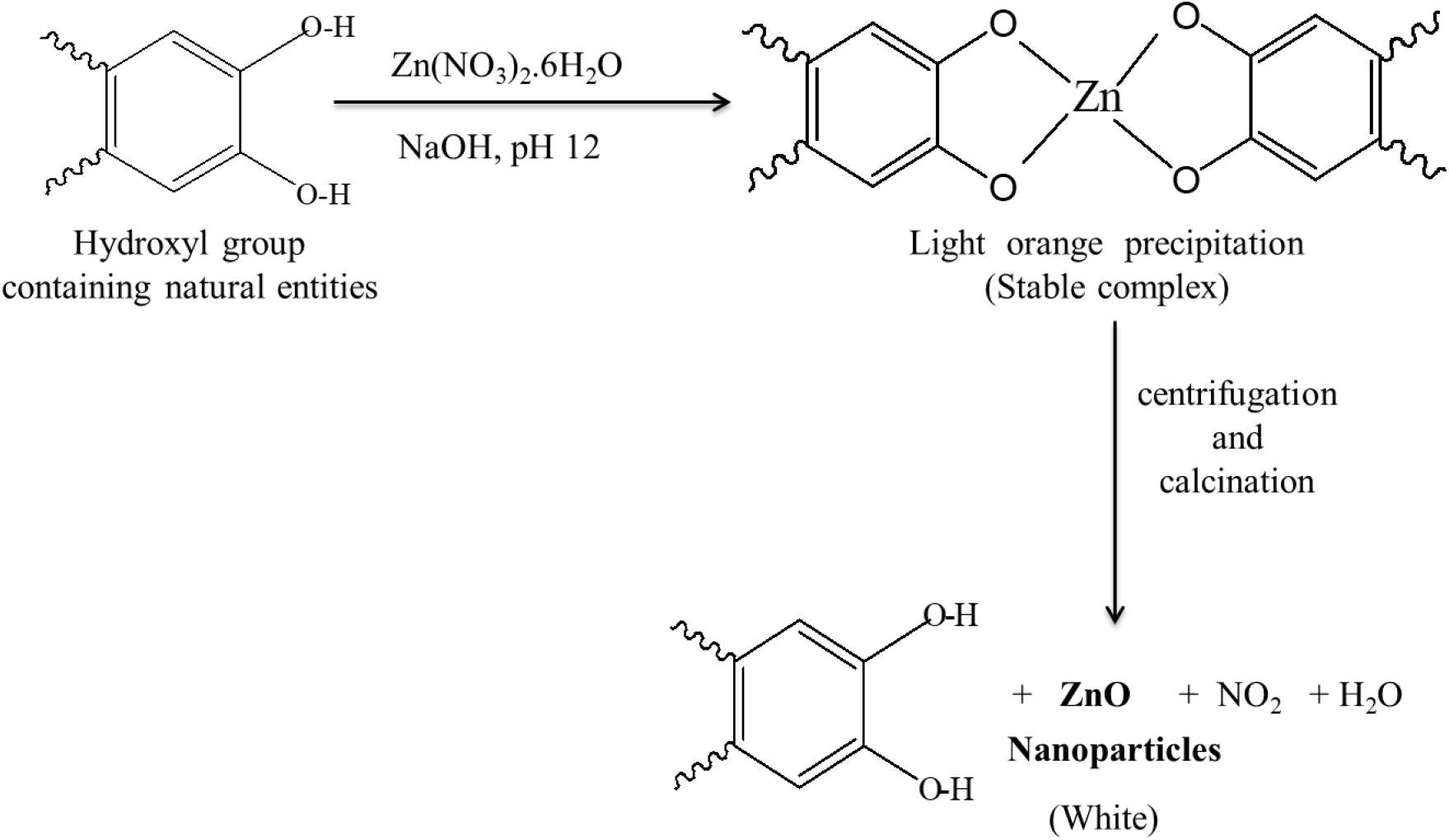
Proposed mechanism of ZnO NPs formation.

### 3.2. Ultraviolet-Visible (UV-Vis) spectroscopy analysis

ZnO NPs generally show UV absorption bands in the λ_max_ ranges from 355 to 380 nm [74, 75]. **Fig. 3** shows the absorption intensity of the prepared ZnO NPs measured in the wavelength range from 300 to 500 nm. The prepared ZnO NPs using *Cocos nucifera* leaf extract showed λ_max_ at 370 nm which is supported by the literature [66–68]. The bandgap energy of ZnO NPs was found to be 3.37 eV as calculated by using Tauc’s plot, which is alike to the reported bandgap energy of ZnO (wide band gap 3.10 – 3.39 eV) [31, 33, 76, 77]. These findings primarily confirm the formation of ZnO NPs following our approaches.

**Fig. 3.**
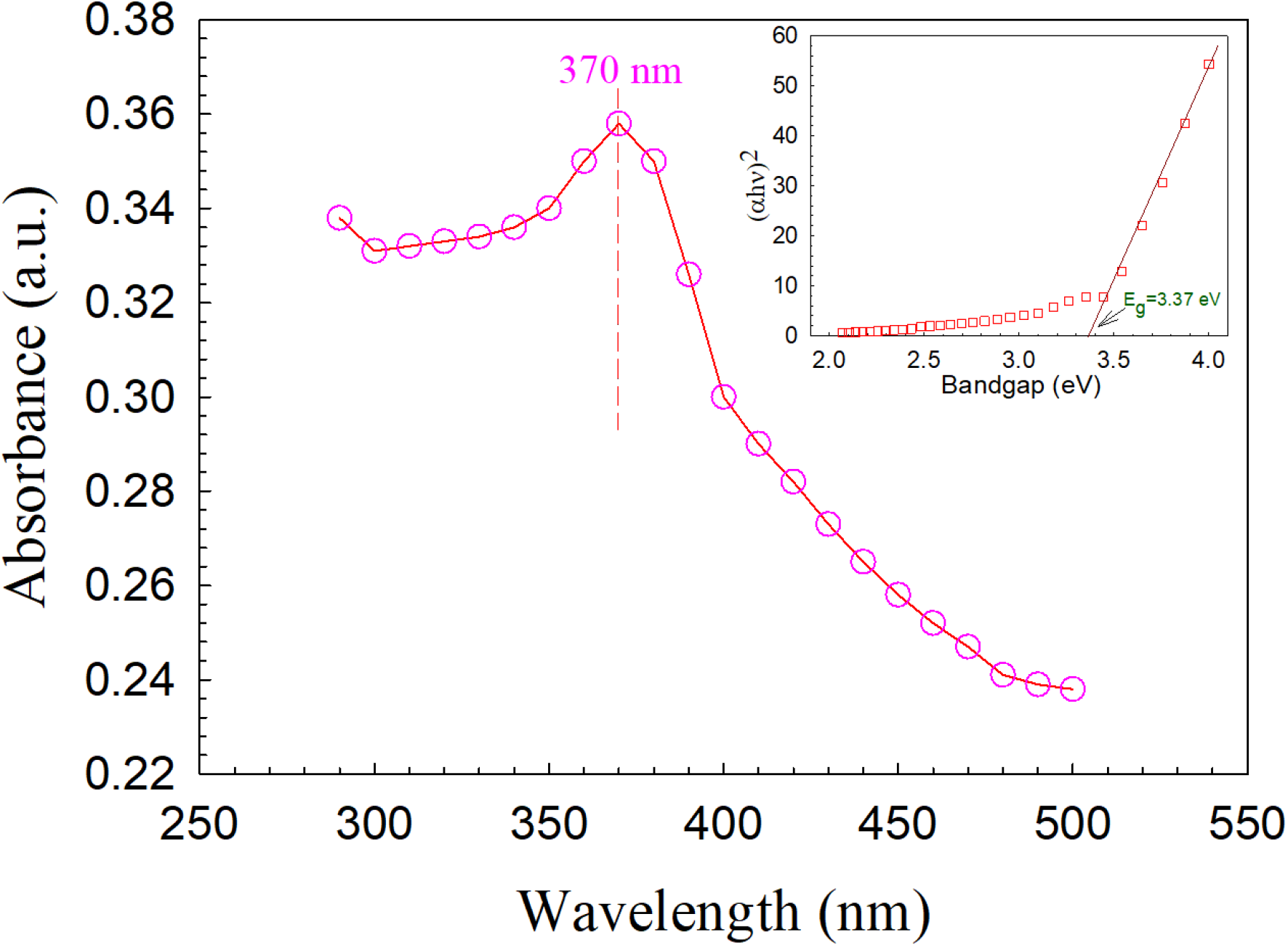
UV–Vis spectrum of ZnO NPs prepared using *Cocos nucifera* leaf extract.

### 3.3. X-ray diffraction analysis

The white powder obtained from the preparation step of the material was subjected to the XRD analysis and the corresponding XRD patterns are shown in **Fig. 4**. The XRD patterns revealed 9 diffraction peaks appearing at 2θ angles of 31.83°, 34.46°, 36.28°, 47.58°, 56.62°, 62.91°, 66.46°, 68.06°, and 69.10° corresponding to the miller indices of 100, 002, 101, 102, 110, 103, 112, 200, and 201, respectively. According to JCPDS card no: 36-1451, the obtained patterns identified our prepared material as ZnO with hexagonal wurtzite structure, space group: P63mc, unit cell volume: 47.62, unit cell parameters: a = b = 3.25 Å and c = 5.21 Å, and α = β = 90° and γ = 120°. The obtained XRD patterns are quite comparable with the previous report [31, 75]. The average crystal size of the prepared ZnO NPs was calculated by using Debye Scherrer’s equation (section 2.4, equation 1) and is found to be 16.6 nm (range:11.9-24.1 nm).

**Fig. 4.**
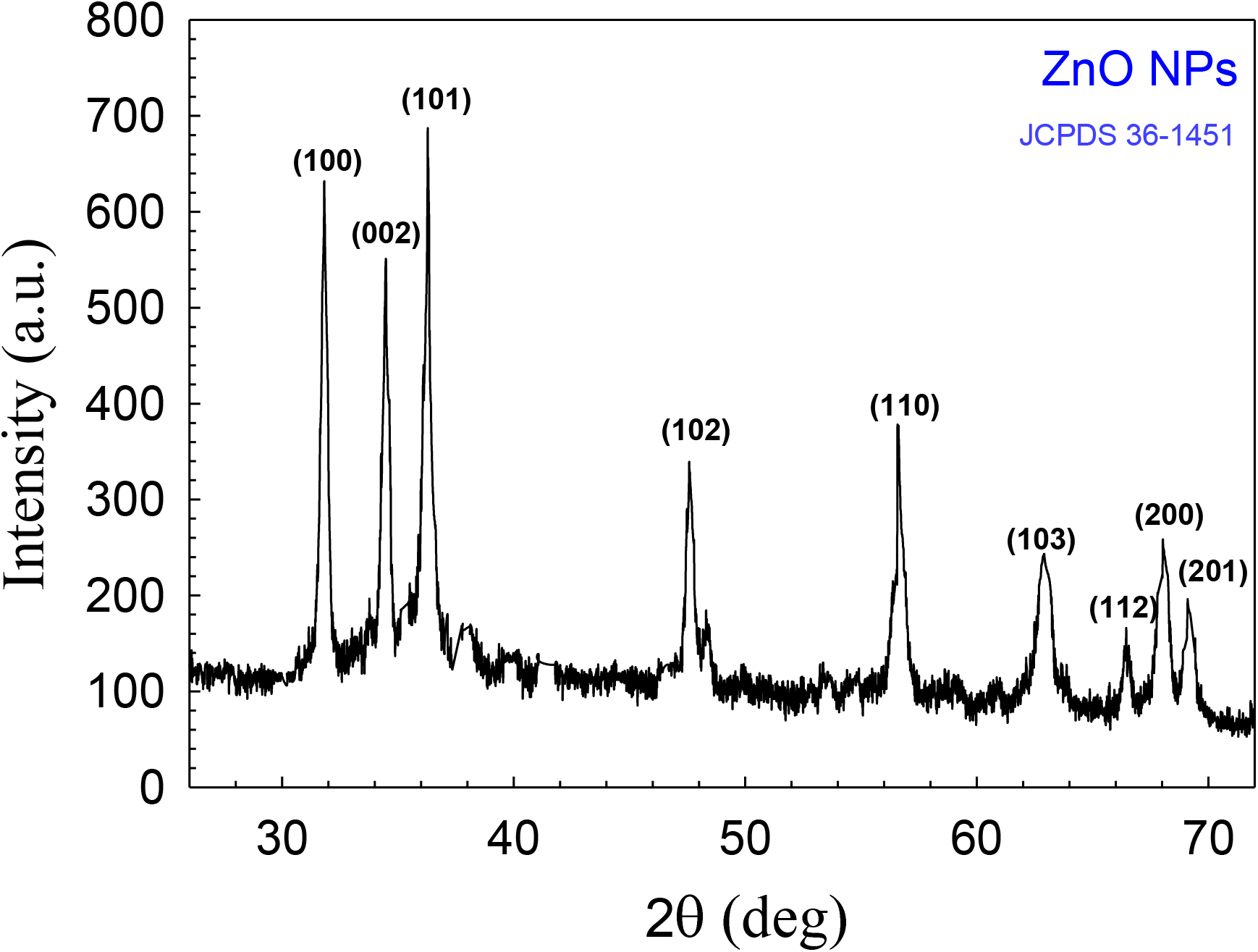
XRD pattern of ZnO Nps prepared using *Cocos nucifera* leaf extract

### 3.4. FTIR spectroscopy analysis

The FTIR spectrum of the prepared ZnO NPs using *Cocos nucifera* leaf extract is illustrated in **Fig. 5**. The inset of Fig. 5 shows the FTIR spectrum of *Cocos nucifera* leaf extract. This spectroscopic measurement was carried out to identify the functional groups of the possible biomolecules responsible for the capping and efficient stabilization of the ZnO NPs. According to the literature [78], the peaks that appeared at (3200-3600) cm^−1^ in the FTIR spectrum can be corroborated by the O-H stretching alcohols, stretching vibrations of the primary and secondary amines, and C−H stretching of alkanes. The peaks observed at 1568, 1411, and 1100 cm^−1^ originated due to the C=C stretching in the aromatic ring in polyphenols and aliphatic amines, while 2300 cm^−1^ originated from di-substituted alkynes, and 550 cm^−1^ from the hexagonal phase of ZnO [78, 79]. As mentioned previously, the *Cocos nucifera* leaf contains alkaloids, steroids, terpenoids, flavonoids, polyphenols, and aromatic hydrocarbons. In this consequence, the results of the FTIR analysis indicate that the functional groups present in the biomolecules of leaf extract as well as phytocompounds such as alkaloids, steroids, terpenoids, flavones, polyphenols, and aromatic hydrocarbons may also act as reducing and capping agent for ZnO NPs formation and prevent agglomeration of the NPs in the aqueous extract medium [47, 80].

**Fig. 5.**
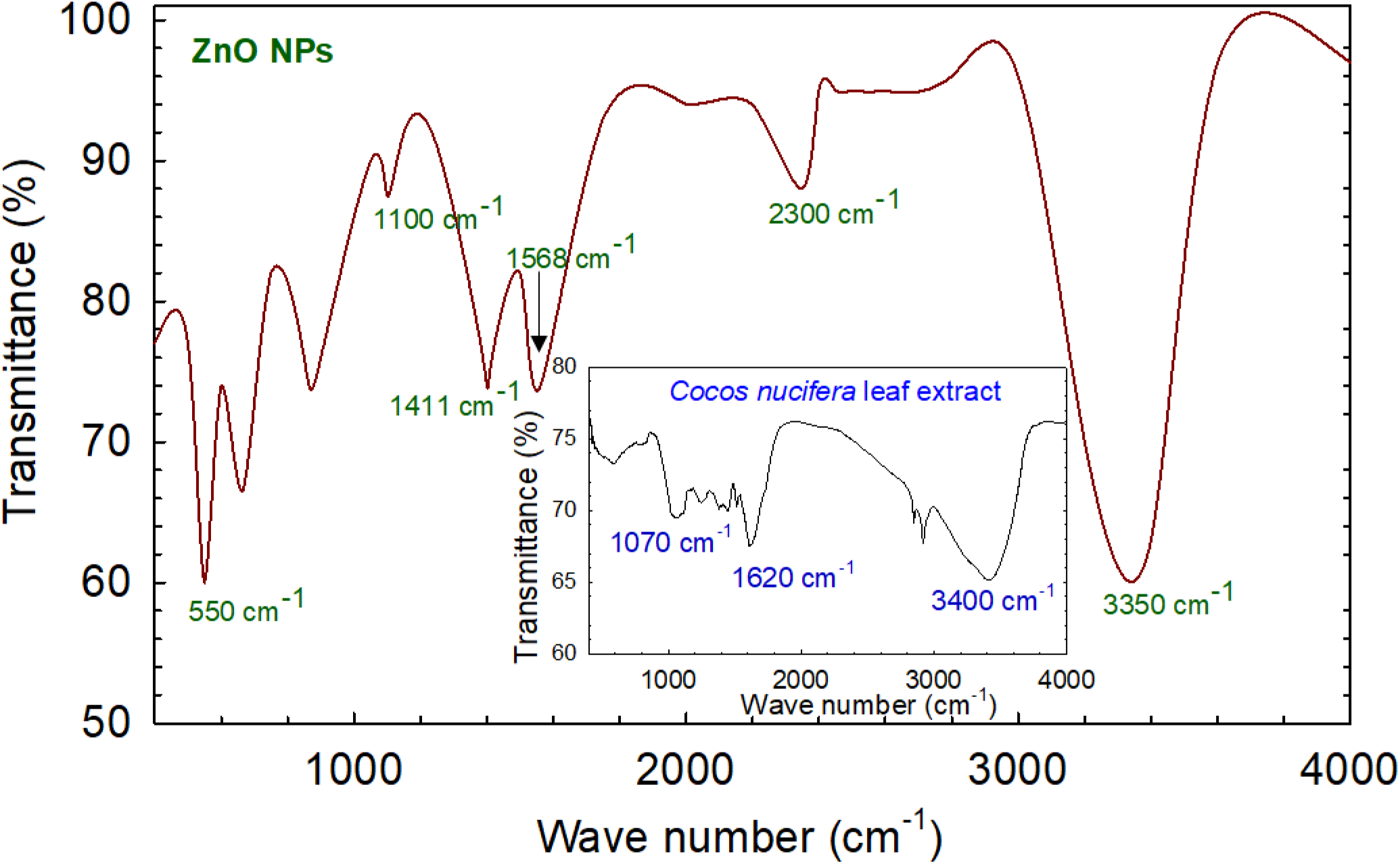
FTIR spectrum of ZnO NPs prepared using *Cocos nucifera* leaf extract. The inset shows the FTIR spectrum of *Cocos nucifera* leaf extract.

### 3.5. Energy dispersive X-ray (EDX) analysis

The elemental composition of ZnO NPs was obtained from EDX analysis. **Fig. 6** shows the existence of chemical elements and their composition in the prepared ZnO NPs. The presence of a large percentage of Zn and O is indicative of ZnO formation. As expected, the atomic percentage of Zn and O are almost equal (ZnO in 1:1 ratio) and then the highest percentage of C along with some other elements such as N, P, S, and Cl originated from the biomolecules of *Cocos nucifera* leaf. The presence of Zn at a high percentage whereas C and other elements in a lower percentage indicate that plant phytochemical groups were involved in reducing and capping the ZnO NPs.

**Fig. 6.**
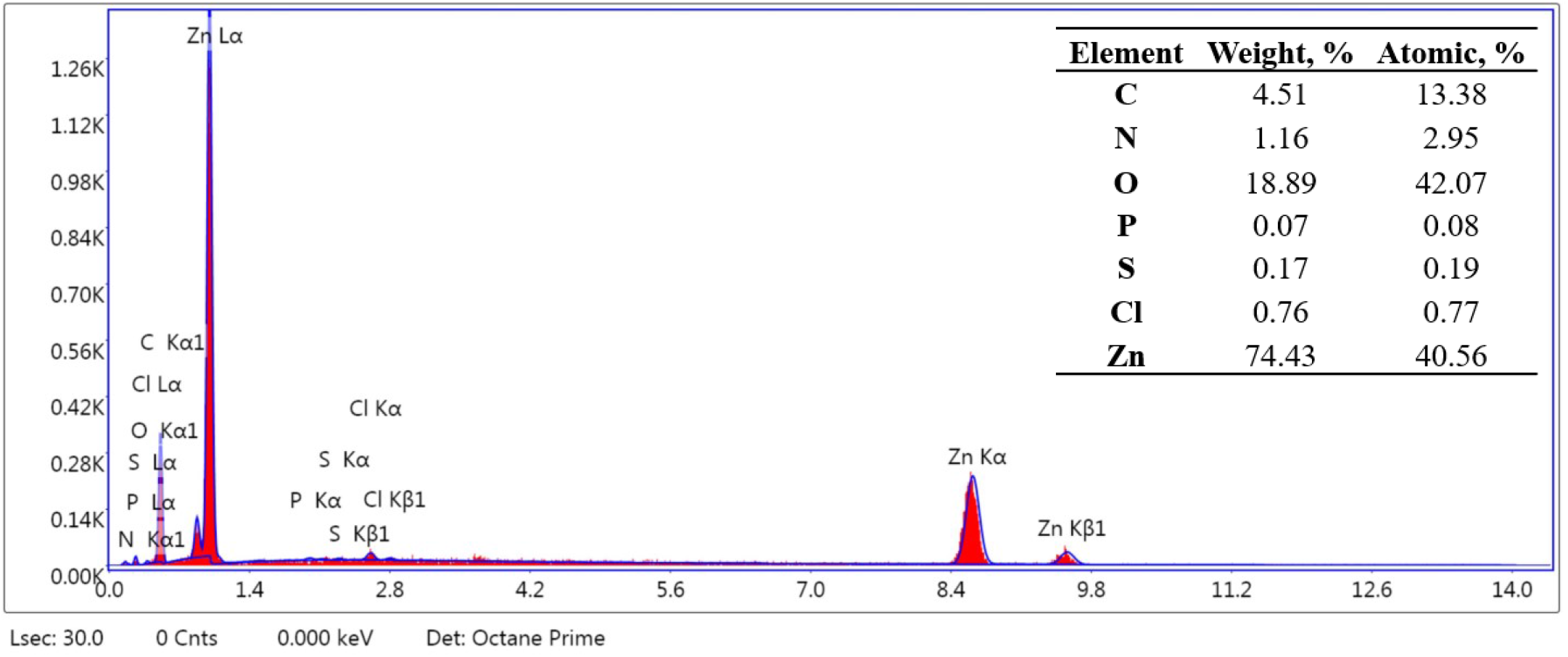
EDX spectrum and elemental composition of ZnO NPs prepared using *Cocos nucifera* leaf extract.

### 3.6. Scanning electron microscopy (SEM) analysis

The surface morphology of the prepared ZnO NPs was studied by SEM analysis. The SEM image of ZnO NPs is depicted in **Fig. 7**, which shows a uniform distribution of flower-shaped ZnO molecules. The calculation of particle size of ZnO NPs from the SEM image using ImageJ software gives the size of NPs as about 15 nm which is agreed with the calculated particle size (16.6 nm) from XRD data. The hydrogen bonding and electrostatic interaction between bioorganic capping molecules and NPs have accumulated them to stay together [81]. Moreover, the SEM image showing ZnO NPs revealed that the NPs are not in direct contact with each other which signifies the stabilization of the NPs by capping agents [82].

**Fig. 7.**
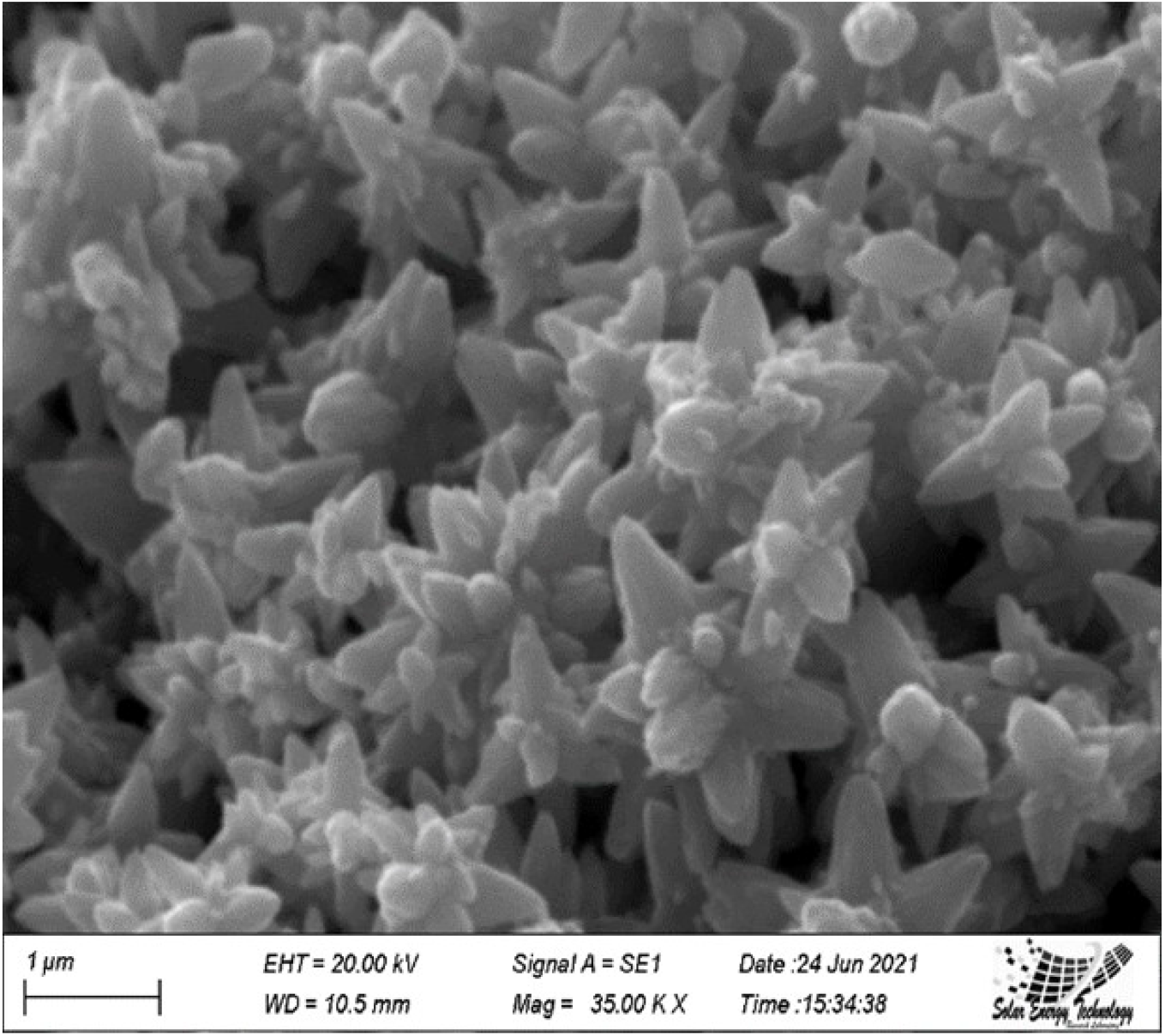
SEM image of ZnO NPs prepared using *Cocos nucifera* leaf extract.

### 3.7. Antimicrobial screening analysis

#### 3.7.1. Mode of action

The ZnO NPs showed good antimicrobial activity against a wide variety of microbes including bacteria and fungi. The prepared NPs interact with the cell membrane of microbes in different pathways such as through the release of ROS (reactive oxygen species), the release of Zn^2+^, and direct contact with the cell membrane. This process damages the cell through DNA disruption, protein denaturation, cellular respiratory disorder, cell membrane damage, and so on. **Fig. 8** illustrates the discussed mechanism which is also proposed in the literature [83, 84].

**Fig. 8.**
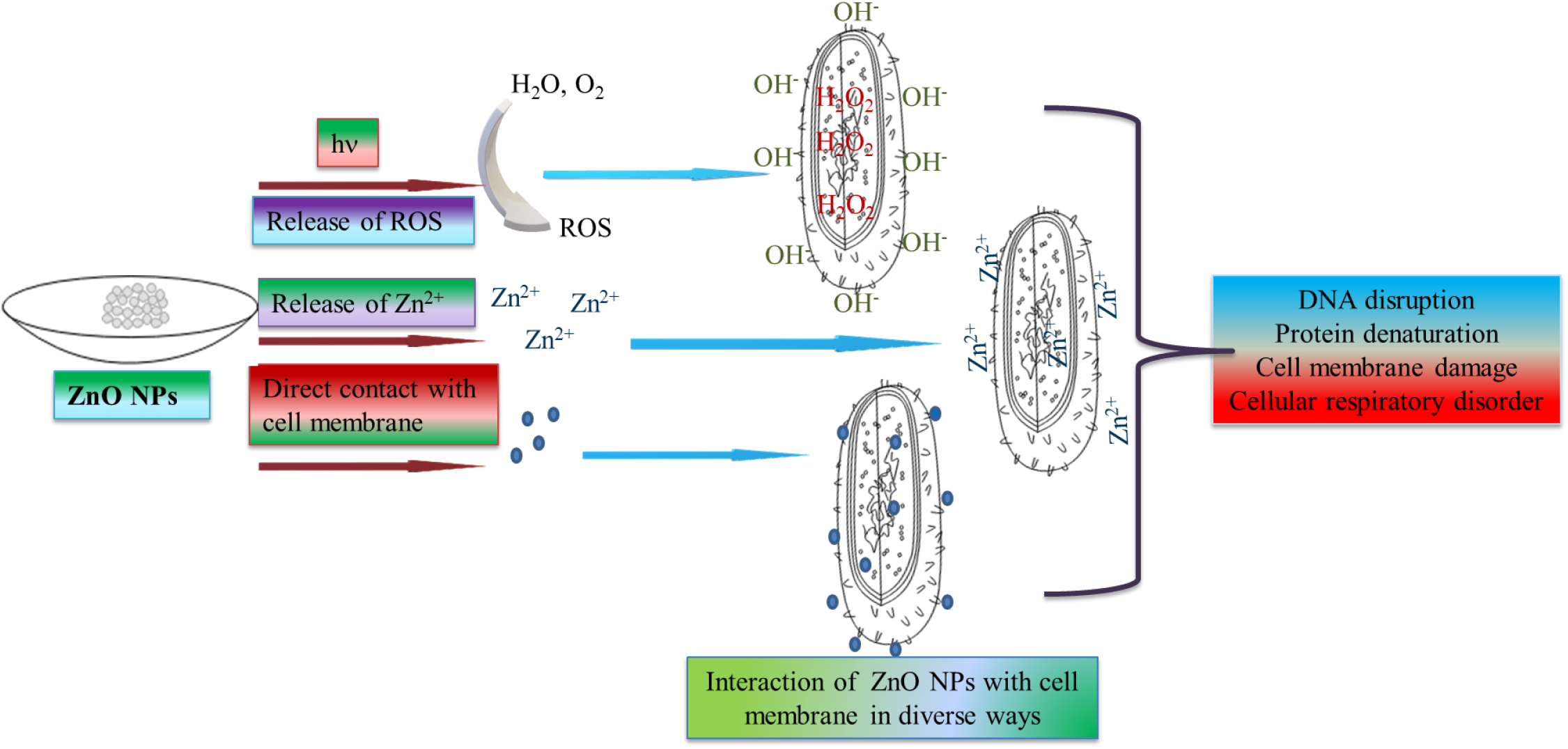
Mode of action of ZnO NPs against microbes (In the figure, ROS means reactive oxygen species).

#### 3.7.2 Antimicrobial activity of ZnO NPs

Antimicrobial activity was studied against three bacterial and two fungal pathogenic strains as shown in **Table 1**. The highest ZOI of ZnO NPs was found to be 14 mm for fungal pathogenic strains of *T. harzianum* which causes infection in renal transplant recipients in humans [85]. Synthesized ZnO nanoparticles show moderate antimicrobial activity (≥10 mm) with gram (+ve) bacteria *S. aureus* (10 mm) and fungus *A. niger* (10 mm) and slight antimicrobial activity (8 – 9 mm) against *B. megaterium* and *S. aureus*.

**Table 1.** ZOI diameters (mm) of ZnO NPs, ceftriaxone, and amphotericin-B against tested bacterial and fungal strains.

| Material | Gram (+ve) Bacteria |  |  | Fungi |  |
| --- | --- | --- | --- | --- | --- |
|  | *B. megaterium* | *S. aureus* | *B. cereus* | *A. niger* | *T. harzianum* |
| ZnO NPs | 08 | 10 | 08 | 10 | 14 |
| Ceftriaxone | 50 | 40 | 20 | - | - |
| Amphotericin-B | - | - | - | 08 | 17 |

#### 3.7.3 Photocatalytic behavior of ZnO NPs

**Fig. 9** (a) demonstrates the photocatalytic behavior of the prepared ZnO NPs which represents the degradation of dye concentration at different time intervals, whereas **9** (b) represents the percentage of dye degradation with respect to time with a clear view (inset) of the color change during the reaction. The initial absorbance of the dye was compared with the final absorption after mixing the prepared ZnO NPs with the dye. It was shown that the absorbance degrades graphically at 640 nm. The mixed solution was discolored after 1 hour and methylene blue dye degraded a maximum of 84.29 % by the ZnO NPs. A possible mechanism is proposed [86, 87] for this degradation process, like the production of ROS (reactive oxygen species) by photo-oxidation (hydroxyl radical generation) and photo-reduction (peroxide radical generation) process [88]. The reaction rate can be calculated using the first-order kinetic equation ln(C_0_/C_t_) = kt, where C_0_ and C_t_ are the initial and final concentration of ZnO NPs and k is the rate constant which is equal to 0.0219 min^−1^, and t is the degradation time showed in **Fig. 9(c)**. Finally, this ROS degraded the dye into mineral acids, CO_2_, and water. The probable mechanism is embedded in the inset of **Fig. 9** (a).

**Fig. 9.**
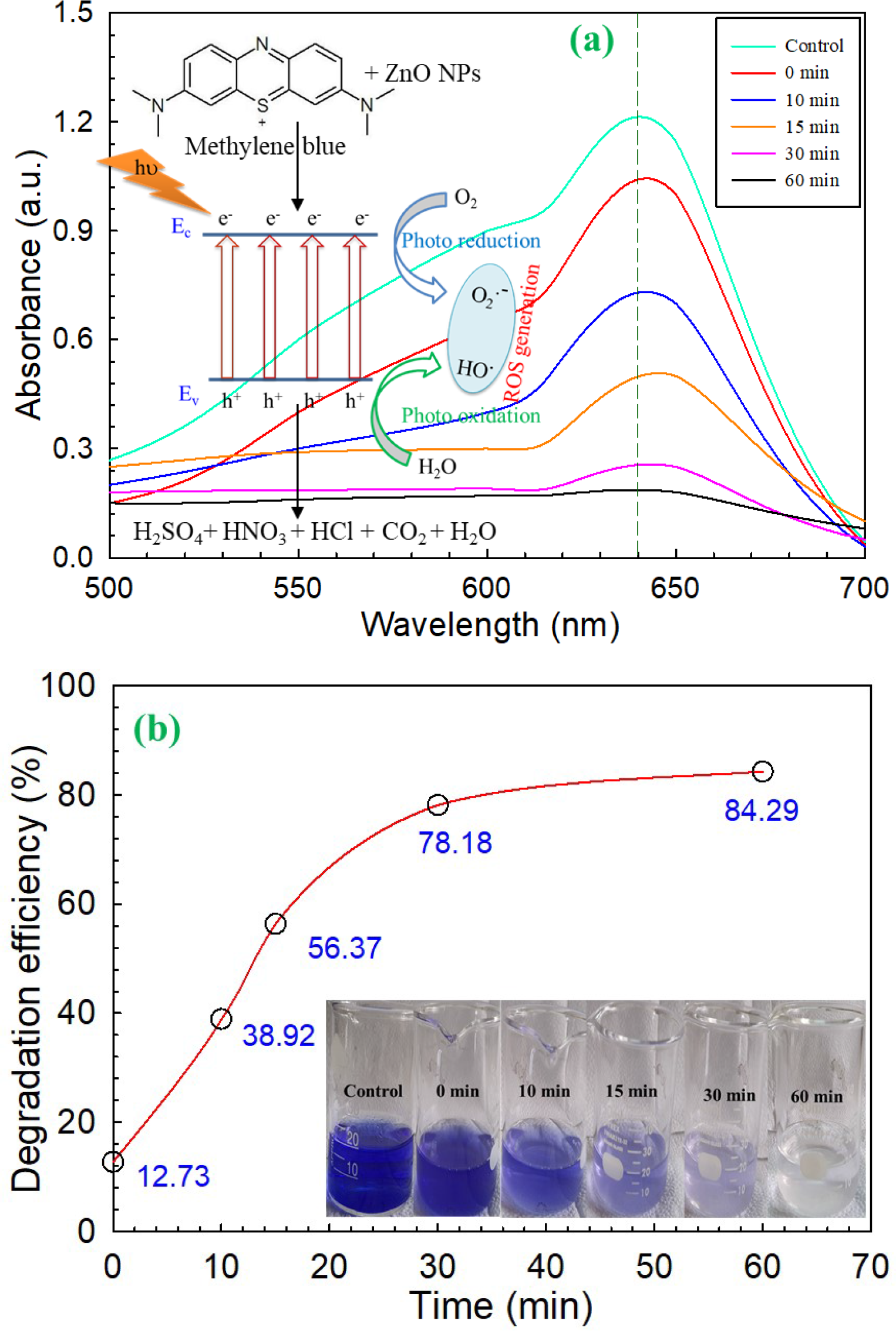
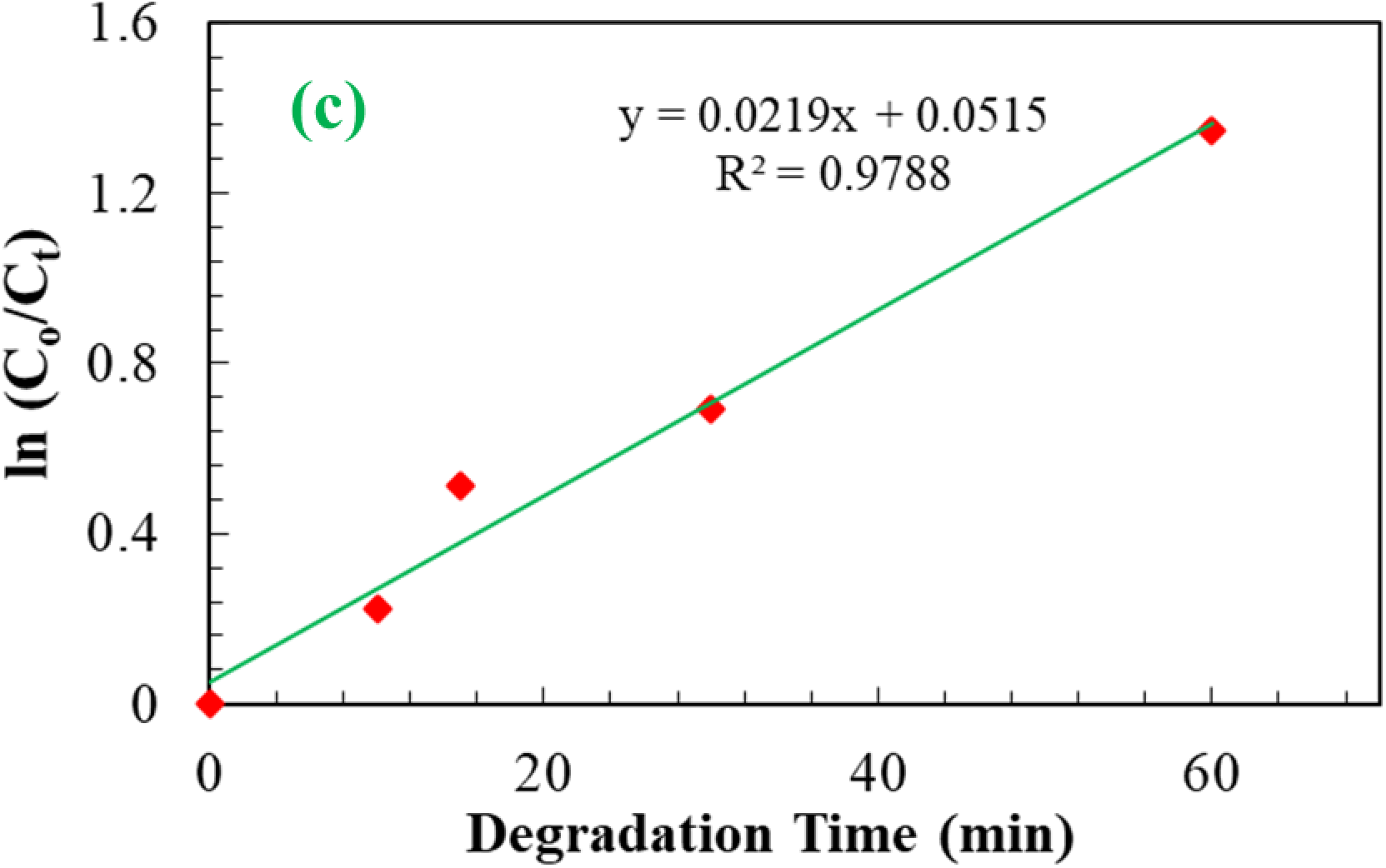
(a) UV–Vis spectra showing the degradation of methylene blue dye through the photocatalysis activity of the prepared ZnO NPs. (b) percentage of degradation of methylene blue dye with respect to a time where the inset figure illustrates the color change with time during the degradation of methyl blue dye with ZnO NPs. (c) shows the reaction kinetics.

#### 3.7.4 Antioxidant activity assay

**Fig. 10** demonstrates the DPPH (2, 2-diphenyl-1-picrylhydrazyl) free radical scavenging activity of the prepared ZnO NPs showing IC_50_ value. The inset of the same figure illustrates the mechanism for DPPH free radical scavenging activity of ZnO NPs following that of Murali et al. [89]. At 517 nm, the absorbance of DPPH decreased with the increase of the concentration of ZnO NPs. This result indicates that the prepared ZnO NPs can inhibit oxidation due to transferring of electron density located at the oxygen atom to the nitrogen atom in DPPH free radical which contains odd electron by n→*π*\* transition [90, 91]. The required concentration of the prepared ZnO NPs to show 50 % inhibition (IC_50_) of DPPH was found to be 764 μg/mL.

**Fig. 10.**
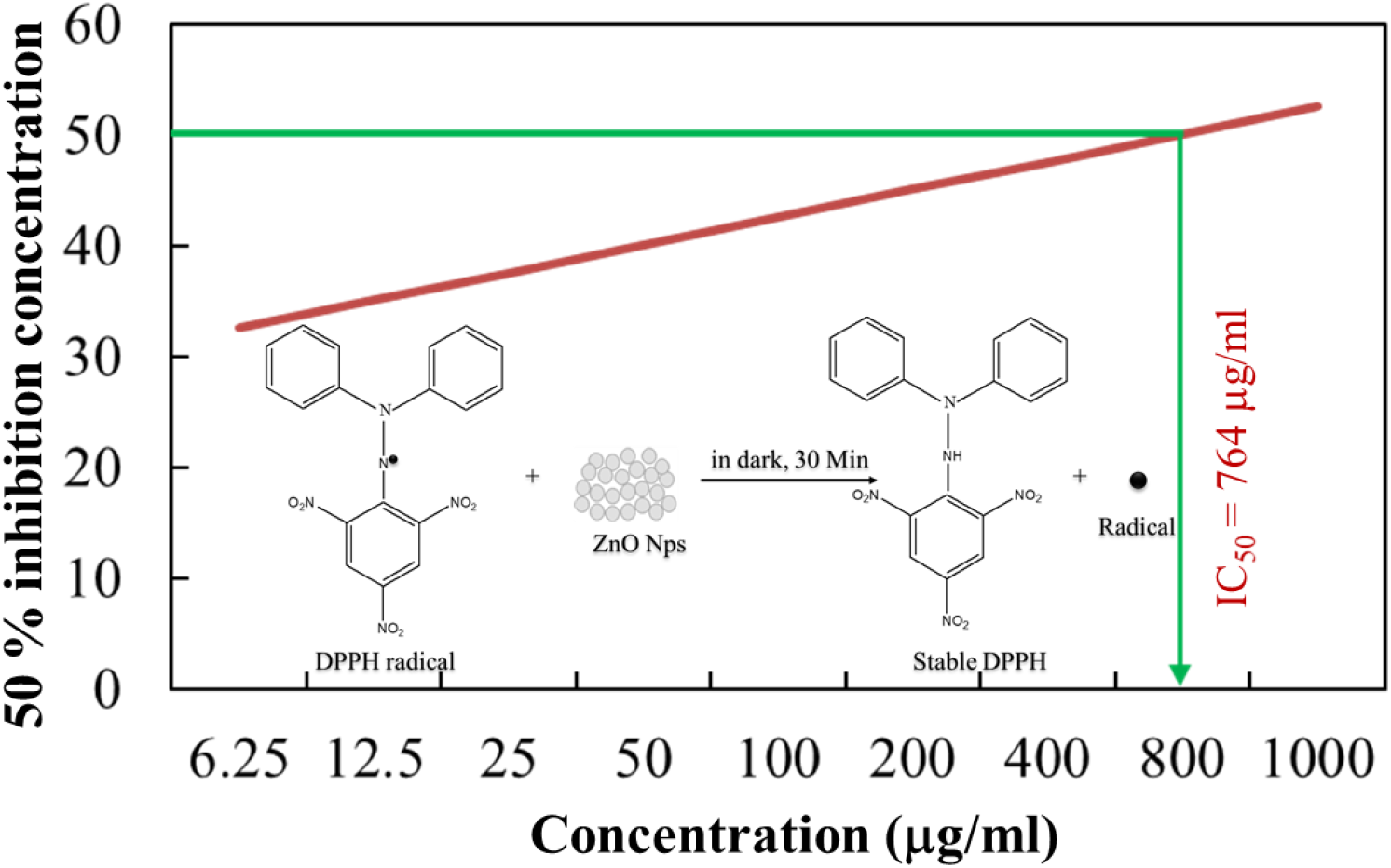
DPPH (2, 2-diphenyl-1-picrylhydrazyl) free radical scavenging activity of the prepared ZnO NPs showing IC_50_ value. Inset shows the mechanism for DPPH free radical scavenging activity of ZnO NPs.

## 4. Conclusions

In this attempt, we have successfully prepared ZnO NPs with an average size of 16.6 nm, using *Cocos nucifera* leaf extract by a facile, inexpensive, and green approach. The prepared NPs were identified and characterized by different techniques such as UV-Vis spectroscopy, XRD, FTIR, EDX, and SEM analyses. The aqueous solution of the prepared ZnO NPs showed absorption maxima, λ_max_ at 370 nm in UV-Vis spectroscopic measurement. The XRD analysis identified the formed ZnO NPs with hexagonal wurtzite structure. The FTIR analysis indicates the presence of some reducing biomolecules associated with some organic functional groups responsible for the encapsulation and stabilization of the NPs. The obtained elemental composition from EDX analysis supports the formation of the desired ZnO NPs. The antimicrobial study of the prepared ZnO NPs showed that the material is very active against various pathogenic bacteria and fungi. The prepared NPs showed high photocatalytic activity while moderate antioxidant activity. Thus, we can conclude that the prepared ZnO NPs could be used in biomedical, medicinal, and pharmaceutical applications; and also, as photocatalysts in the dye degradation process.

## Acknowledgment

The authors highly acknowledge the Organic Chemistry Research Laboratory, Department of Chemistry, Comilla University, Cumilla for the laboratory and instrumental facilities. We also acknowledge the Bangladesh Council of Scientific and Industrial Research (BCSIR) and Bangladesh Atomic Energy Commission for SEM-EDX and XRD analysis. The authors are thankful to the Department of Pharmacy, Comilla University, and the Department of Chemistry, Jagannath University, Dhaka for allowing us to experiment on the antioxidant and antimicrobial study.

## Ethical Statements

The study was conducted according to the guidelines of the planning and development (P&D) committee of the Department of Chemistry, Comilla University (Ref: Chem/P&D/100/10-15;14/09/2020). *Cocos nucifera* leaves were used in this study which was kindly provided by Dr. Mohammad Anowar Hossain Bhuyan, Joint Director, Bangladesh Academy for Rural Development (BARD), Cumilla 3503, Bangladesh, in September 2020, and were authenticated by Professor Dr. Shaikh Bokhtear Uddin, Department of Botany, University of Chittagong, Chattogram-4331, Bangladesh. A voucher specimen of this species was deposited to the Chittagong University Herbarium with Accession number (SBU 210222-32660 CUH).

## Availability of Data

Our data are deposited at Dryad as (DOI): https://doi.org/10.5061/dryad.tht76hf27.

## Author Contribution

***Farjana Rahman**:* Experiment, Investigation, Methodology, Data curation, Formal analysis, Writing – original draft.

***Md Abdul Majed Patwary**:* Supervision, Funding acquisition, Experiment, Investigation, Methodology, Data curation, Formal analysis, Writing – original draft, review & editing.

***Md. Abu Bakar Siddique**:* Experiment, Formal analysis, Writing – review & editing. ***Muhammad Shahriar Bashar**:* Experiment, Formal analysis, Writing – review & editing. ***Md. Aminul Haque***: Experiment, Formal analysis, Writing – review & editing.

***Beauty Akkter**:* Experiment, Investigation, Methodology, Data curation.

***Rimi Rashid**:* Experiment, Formal analysis, Writing – review & editing.

***Md. Anamul Haque**:* Experiment, Formal analysis, Writing – review & editing.

***A. K. M. Royhan Uddin**:* Supervision, Funding acquisition, Conceptualization, Methodology, Formal analysis, Writing – review & editing.

## References

1. A. Rastogi, M. Zivcak, O. Sytar, H. M. Kalaji, X. He, S. Mbarki and M. Brestic, 2017, Impact of metal and metal oxide nanoparticles on plant: A critical review. Front. Chem., 5, 1–16. doi: 10.3389/fchem.2017.00078

2. V. Marassi, L. D. Cristo, S. G. J. Smith, S. Ortelli, M. Blosi, A. L. Costa, P. Reschiglian, Y. Volkov and A. Prina-Mello, 2018, Silver nanoparticles as a medical device in healthcare settings: a five-step approach for candidate screening of coating agents. R. Soc. open sci. 5: 171113. 171113. DOI: https://doi.org/10.1098/rsos.171113

3. A. M. Awwad, M. W. Amer, N. M. Salem and A. O. Abdeen, 2020, Green synthesis of zinc oxide nanoparticles (ZnO-NPs) using Ailanthus altissima fruit extracts and antibacterial activity. Chem. Int., 6, 151–159. https://doi.org/10.5281/zenodo.3559520

4. O. V. Kharissova, B. I. Kharisov, C. M. O. González, Y. P. Méndez and I. López, 2019, Greener synthesis of chemical compounds and materials. R. Soc. open sci. 6,191378. 191378. DOI: https://doi.org/10.1098/rsos.191378

5. M. Anbuvannan, M. Ramesh, G. Viruthagiri, N. Shanmugam and N. Kannadasan, 2015, Synthesis, characterization and photocatalytic activity of ZnO nanoparticles prepared by biological method. Spectrochim. Acta A Mol. Biomol. Spectrosc., 143, 304–308. http://dx.doi.org/10.1016/j.saa.2015.01.124

6. B. C. Ghos, S. F. U. Farhad, M. A. M. Patwary, S. Majumder, M. A. Hossain, N. I. Tanvir, M. A. Rahman, T. Tanaka and Q. Guo, 2021, Influence of the substrate, process conditions, and postannealing temperature on the properties of ZnO thin films grown by the successive ionic layer adsorption and reaction method. J. ACS Omega, 6, 2665–2674. https://dx.doi.org/10.1021/acsomega.0c04837

7. M. Sundrarajan, S. Ambika and K. Bharathi, 2015, Plant-extract mediated synthesis of ZnO nanoparticles using Pongamia pinnata and their activity against pathogenic bacteria. Adv. Powder Technol., 26, 1294–1299. http://dx.doi.org/10.1016/j.apt.2015.07.001

8. W. Beek, M. Wienk and R. Janssen, 2004, Efficient hybrid solar cells from ZnO nanoparticles and a conjugated polymers. Adv. Mater., 16, 1009–13. doi:10.1002/adma.200306659

9. A. Suliman, Y. Tang and L. Xu, 2007, Preparation of ZnO nanoparticles and nanosheets and their application to dye-sensitized solar cells. Sol. Energy Mater. Sol. Cell, 91, 1658–1662. doi:10.1016/j.solmat.2007.05.014

10. S. Majumder, N. I. Tanvir, B. C. Ghos, M. A. M. Patwary, M. A. Rahman, M.A. Hossain and S. F. U. Farhad, 2020, Optimization of the growth conditions of Cu_2_O thin films and subsequent fabrication of Cu_2_O/ZnO heterojunction by m-SILAR method. IEEE WIECON-ECE, 139–142. doi: 10.1109/WIECON-ECE52138.2020.9397989

11. T. Li, N. Bao, A. Geng, H. Yu, Y. Yang and X. Dong, 2018, Study on room temperature gas-sensing performance of CuO film-decorated ordered porous ZnO composite by In_2_O_3_ sensitization. R. Soc. open sci., 5: 171788. 171788. DOI: https://doi.org/10.1098/rsos.171788

12. B. Baruwati, D. Kumar and S. Manorama, 2006, Hydrothermal synthesis of highly crystalline ZnO nanoparticles: A competitive sensor for LPG and EtOH. Sen. Actu. B. Chem., 119, 676–82. doi:10.1016/j.snb.2006.01.028

13. M. Vaseem, A. Umar and Y. Hahn, 2010, ZnO nanoparticles: growth, properties, and applications. Metal Oxide Nanostructures and Their Applications, ed. A. Umar and Y. B. Hahn, American Scientific Publishers, USA, 5, ch. 4, pp. 1–36.

14. S. Chang and K. Chen, 2012, Zinc Oxide nanoparticle photodetector. J Nanomater, 2012, 1–5. doi:10.1155/2012/602398

15. . T. Mishchenko, E. Mitroshina, I. Balalaeva, O. Krysko, M. Vedunova, D. Krysko, 2019, An emerging role for nanomaterials in increasing immunogenicity of cancer cell death. Biochim Biophys Acta Rev Canc, 1871, 99–108. doi:10.1016/j.bbcan.2018.11.004

16. R. Dadi, R. Azouani, M. Traore, C. Mielcarek and A. Kanaev, 2019, Antibacterial activity of ZnO and CuO nanoparticles against gram positive and gram negative strains. Mater Sci Eng C Mater Biol Appl. doi:10.1016/j.msec.2019.109968

17. R. D. Umrani and K. M. Paknikar, 2014, Zinc oxide nanoparticles show antidiabetic activity in streptozotocin-induced Types 1 and 2 diabetic rats. Nanomedicine, 9, 89–104. doi:10.2217/nnm.12.205

18. R. M. El-Gharbawy, A. M. Emara and S. E. Abu-Risha, 2016, Zinc oxide nanoparticles and a standard antidiabetic drug restore the function and structure of beta cells in Type-2 diabetes. Biomed Pharmacother, 84, 810–820. doi:10.1016/j.biopha.2016.09.068

19. Y. Wang, S. Song, J. Liu, D. Liu and H. Zhang, 2015, ZnO-functionalized upconverting nanotheranostic agent: multi-modality imaging-guided chemotherapy with on-demand drug release triggered by pH. Angew Chem Int Ed Engl, 54, 536–40. doi:10.1002/anie.201409519

20. J. E. Eixenberger, C.B. Anders, K. Wada, K. M. Reddy, R. J. Brown, J. Moreno-Ramirez, A. Weltner, K. Chinnathambi, D. Tenne, D. Fologea and D. Wingett, 2019, Defect engineering of ZnO nanoparticles for bio-imaging applications. ACS Appl Mater Interfaces, 11, 24933–44. doi:10.1021/acsami.9b01582

21. A. Hussain, M. Oves, M. F. Alajmi, I. Hussain, S. Amir, J. Ahmed, M. T. Rehman, H. R. El-Seedif and I. Ali, 2019, Biogenesis of ZnO nanoparticles using *Pandanus odorifer* leaf extract: anticancer and antimicrobial activities. RSC Advances, 9, 15357–15369. doi:10.1039/c9ra01659g

22. S. Shahid, U. Fatima, R. Sajjad, S. A. Khan, 2019, Bioispired nanotheranostic agent: Zinc Oxide; green synthesis and biomedical potential. Dig. Jour. of Nanomat. and Biostruc., 14, 1023–1031.

23. M. Batool, S. Khurshid, W. M. Daoush, S.A. Siddique and T. Nadeem, 2021. Green synthesis and biomedical applications of ZnO nanoparticles: role of PEGylated-ZnO nanoparticles as doxorubicin drug carrier against MDA-MB-231(TNBC) cells line. Crystals, 11, 1–19. https://doi.org/10.3390/cryst11040344

24. N. Elavarasan, K. Kokila, G. Inbasekar and V. Sujatha, 2016, Evaluation of photocatalytic activity, antibacterial and cytotoxic effects of green synthesized ZnO nanoparticles by *Sechium edule* leaf extract Res Chem Intermed, 43, 3361–3376.

25. A. H. Aalami, M. Mesgari and A. Sahebkar, 2020, Synthesis and characterization of green Zinc Oxide nanoparticles with antiproliferative effects through apoptosis induction and microRNA modulation in breast cancer cells. Bioinorg Chem Appl, 2020, 1–17. https://doi.org/10.1155/2020/8817110

26. R. L. Kalyani, S. V. N. Pammi, P. P. N. V. Kumar, P. V. Swamy, K. V. R. Murthy, 2019, Antibiotic potentiation and anti-cancer competence through bio-mediated ZnO nanoparticles. Mater Sci Eng C Mater Biol Appl, 103. DOI: https://doi.org/10.1016/j.msec.2019.109756

27. M. Shahnaza, M. Danish, M. Hisyamuddin, B. Ismail, M. T. Ansari and M. N. M. Ibrahim, 2019, Anticancer and apoptotic activity of biologically synthesized zinc oxide nanoparticles against human colon cancer HCT-116 cell line-in vitro study. Sust. Chem. and Pharm., 14, 1–7. DOI: https://doi.org/10.1016/j.scp.2019.100179

28. S. Rajeshkumar, S. V. Kumar, A. Ramaiah, H. Agarwal, T. Lakshmi and S. M. Roopan, 2018, Biosynthesis of zinc oxide nanoparticles using *Mangifera indica* leaves and evaluation of their antioxidant and cytotoxic properties in lung cancer (A549) cells. Enzyme Microb Technol, 117, 91–95. DOI: https://doi.org/10.1016/j.enzmictec.2018.06.009

29. S. Vijayakumar, B. Vaseeharan, B. Malaikozhundan and M. Shobiya, *Biomed Pharmacother*, 2016, *Laurus nobilis* leaf extract mediated green synthesis of ZnO nanoparticles: Characterization and biomedical application. 84, 1213–1222. DOI: http://dx.doi.org/10.1016/j.biopha.2016.10.038

30. R. Dobrucka, and J. Dlugaszewska, 2016, Biosynthesis and antibacterial activity of Zno nanoparticles using *Trifolium pratense* flower extract. Saudi J Biol Sci, 23, 517–23. DOI: http://dx.doi.org/10.1016/j.sjbs.2015.05.016

31. S. Azizi, R. Mohamad, A. Bahadoran, S. Bayat, R. A. Rahim, A. Ariff and W. Z. Saad, 2016, Effect of annealing temperature on antimicrobial and structural properties of bio-synthesized Zinc Oxide nanoparticles using flower extract of *Anchusa italic*. J Photochem Photobiol B, 161, 441–449. doi: 10.1016/j.jphotobiol.2016.06.007

32. U. L. Ifeanyichukwu, O. E. Fayemi and C. N. Ateba, 2020, Green synthesis of zinc oxide nanoparticles from Pomegranate (*Punica granatum*) extracts and characterization of their antibacterial activity. Molecules, 2020. 25, 1–22. DOI: 10.3390/molecules25194521

33. E. Zare, S. Pourseyedi, M. Khatami and E. Darezereshki, 2017, Simple biosynthesis of zinc oxide nanoparticles using nature’s source, and it’s in vitro bio-activity. Jour. of Mol. Struct., 1146, 96–103. DOI: 10.1016/j.molstruc.2017.05.118

34. B. Malaikozhundan, B. Vaseeharan, S. Vijayakumar, K. Pandiselvi, R. Kalanjiam, K. Murugan, and G. Benelli, 2017, Biological therapeutics of *Pongamia pinnata* coated zinc oxide nanoparticles against clinically important pathogenic bacteria, fungi and MCF-7 breast cancer cells. Microb Pathog, 104, 68–277. DOI: 10.1016/j.micpath.2017.01.029

35. V. Anbukkarasi, R. Srinivasan and N. Elangovan, 2015, Antimicrobial activity of green synthesized zinc oxide nanoparticles from *Emblica officinalis*. Int. J. Pharm. Sci. Rev. Res., 33, 110–115.

36. K. Vimalaa, S. Sundarraja, M. Paulpandia, S. Vengatesanc and S. Kannan, 2014, Green synthesized doxorubicin loaded zinc oxide nanoparticles regulates the Bax and Bcl-2 expression in breast and colon carcinoma. J. Proc. Biochem., 49, 160–172. DOI: http://dx.doi.org/10.1016/j.procbio.2013.10.007

37. R. Anitha, K. V. Ramesh, T. N. Ravishankar, K. H. S. Kumar and T. Ramakrishnappa, 2018, Cytotoxicity, antibacterial and antifungal activities of ZnO nanoparticles prepared by *Artocarpus gomezianus* fruit mediated facile green combustion method. J of Sci. Adv. Mat. and Dev., 3, 440–451. DOI: https://doi.org/10.1016/j.jsamd.2018.11.001

38. N. K. Rajendran, B. P. George, N. N. Houreld and H. Abrahamse, 2021, Synthesis of zinc oxide nanoparticles using *Rubus fairholmianus* root extract and their activity against pathogenic bacteria. molecules, 26, 1–11. DOI: https://doi.org/10.3390/molecules26103029

39. K. S. Prasad, S. K. Prasad, R. Veerapur, G. Lamraoui, A. Prasad, M. N. N. Prasad, S. K. Singh, N. Marraiki, A. Syed and C. Shivamallu, 2021, Antitumor potential of green synthesized ZnONPs using root extract of *Withania somnifera* against human breast cancer cell line. separations, 8, 1–9. DOI: https://doi.org/10.3390/separations8010008

40. A. C. Janaki, E. Sailatha and S. Gunasekaran, 2015, Synthesis, characteristics and antimicrobial activity of ZnO nanoparticles. Spectro. Acta Part A: Mol. and Biomol. Spectro., 144, 17–22. doi: http://dx.doi.org/10.1016/j.saa.2015.02.041

41. K. Dulta, G. K. Ağçeli, P. Chauhan, R. Jasrotia and P. K. Chauhan, 2020, A novel approach of synthesis zinc oxide nanoparticles by *Bergenia ciliata* rhizome extract: antibacterial and anticancer potential. J. Inorg. and Org. met. Poly. and Mater., 31, 180–190. DOI: https://doi.org/10.1007/s10904-020-01684-6

42. C. Joel and M. S. M. Badhusha, 2016, Green synthesis of ZnO Nanoparticles using *Phyllanthus embilica* stem extract and their antibacterial activity. Der Pharmacia Lettre, 8, 218–223.

43. M. A. Ansari, M. Murali, D. Prasad, M. A. Alzohairy, A. Almatroudi, M. N. Alomary, A. C. Udayashankar, S. B. Singh, S. M. M. Asiri, B. S. Ashwini, H. G. Gowtham, N. Kalegowda, K. N. Amruthesh, T. R. Lakshmeesha, and S. R. Niranjana, 2020, *Cinnamomum verum* bark extract mediated green synthesis of ZnO nanoparticles and their antibacterial potentiality. Biomolecules, 10, 1–15. doi:10.3390/biom10020336

44. H. Umar, D. Kavaz and N. Rizaner, 2019, Biosynthesis of zinc oxide nanoparticles using *Albizia lebbeck* stem bark, and evaluation of its antimicrobial, antioxidant, and cytotoxic activities on human breast cancer cell lines. Int J Nanomedicine, 14, 87–100.

45. S. N. A. M. Sukri, K. Shameli, M. Mei-T. Wong, S. Y. Teow, J. Chew and N. A. Ismail, 2019, Cytotoxicity and antibacterial activities of plant-mediated synthesized zinc oxide (ZnO) nanoparticles using *Punica granatum* (pomegranate) fruit peels extract. J of Mol. Struc., 1189, 57–65. DOI: https://doi.org/10.1016/j.molstruc.2019.04.026

46. J. Ruangtong, J. T-Thienprasert, and N. P. T-Thienprasert, 2020, Green synthesized ZnO nanosheets from banana peel extract possess anti-bacterial activity and anti-cancer activity. Materials Today Communications, 24. doi: https://doi.org/10.1016/j.mtcomm.2020.101224

47. A.K.M.R. Uddin, M.A.B. Siddique, F. Rahman, A. K. M. A. Ullah and R. Khan, 2020, *Cocos nucifera* leaf extract mediated green synthesis of silver nanoparticles for enhanced antibacterial Activity. J Inorg Organomet Polym, 30, 3305–3316. DOI: https://doi.org/10.1007/s10904-020-01506-9

48. T. Pradeepkumar, B. Sumajyothibhaskar, K. N. Satheesan, 2008, Horticulture Science Series: Vol: XI: Management of Horticultural Crops (2 Parts-Set). New India Publishing, 11, 539–587.

49. An excellent online database of a huge range of trees giving very good information on each plant – its uses, ecology, identity, propagation, pests etc., World Agroforesty Centre, http://www.worldagroforestry.org/

50. R. A. DeFilipps, S. L. Maina and J. Crepin, An excellent guide to the traditional medicinal uses of the plants of the region, Medicinal Plants of the Guianas, Website: http://botany.si.edu/bdg/medicinal/index.html

51. O. A. Akinyemi and F. S. Oyelere, phytochemical profile of selected morphological organs of *Cocos Nucifera L.*, ejbps, 2019, 6, 11, 54–58.

52. E. Zare, S. Pourseyedi, M. Khatami, E. Darezereshki, 2017, Simple biosynthesis of zinc oxide nanoparticles using nature’s source, and it’s in vitro bio-activity. J of Mol. Struc., 1146, 96–103. DOI: 10.1016/j.molstruc.2017.05.118

53. P. K. Ghosh, P. Bhattacharjee, S. Mitra, and M. P. Sarkar, 2014, Physicochemical and Phytochemical Analyses of Copra and Oil of *Cocos nucifera L.* (West Coast Tall Variety). International journal of food science, 2014. DOI: https://doi.org/10.1155/2014/310852

54. S. M. Roopan, 2016, An overview of phytoconstituents, biotechnological applications, and nutritive aspects of coconut *(Cocos nucifera)*. Appl Biochem Biotechnol, 179, 1309–1324. DOI: https://doi.org/10.1007/s12010-016-2067-y

55. M. Satheshkumar, B. Anand, A. Muthuvel, M. Rajarajan, V. Mohana, A. Sundaramanickam, 2020, Enhanced photocatalytic dye degradation and antibacterial activity of biosynthesized ZnO‑NPs using curry leaves extract with coconut water. Nanotechnol. Environ. Eng., 5, 1–11. DOI: https://doi.org/10.1007/s41204-020-00093-x

56. P. I. Priyatharesini, R. Ganesamoorthy and R. Sudha, 2020, Synthesis of zinc oxide nanoparticle using *Cocos nucifera* male flower extract and analysis their antimicrobial Activity. J of Pharm and Tech, 13, 2151–2154. DOI: http://dx.doi.org/10.5958/0974-360X.2020.00386.8

57. A. N. D. Krupa and R. Vimala, 2016, Evaluation of tetraethoxysilane (TEOS) sol–gel coatings, modified with green synthesized zinc oxide nanoparticles for combating microfouling. Mater Sci Eng C Mater Biol Appl., 61, 728–735. doi: 10.1016/j.msec.2016.01.013.

58. E. D. Mahmoud, K. M. Al‑Qahtani, S. O. Alflaij, S. F. Al‑Qahtani, F. A. Alsamhan, Green copper oxide nanoparticles for lead, nickel, and cadmium removal from contaminated water. Scientific Reports, (2021) 11:12547. https://doi.org/10.1038/s41598-021-91093-7

59. J. Kadam, S. Madiwale, B. Bashte, S. Dindorkar, P. Dhawal, P. More, Green mediated synthesis of palladium nanoparticles using aqueous leaf extract of *Gymnema sylvestre* for catalytic reduction of Cr (VI), SN Applied Sciences (2020) 2:1854. https://doi.org/10.1007/s42452-020-03663-5

60. K. V. Alex, P. T. Pavai, R. Rugmini, M. S. Prasad, K. Kamakshi, and K. Chandra Sekhar, Green Synthesized Ag Nanoparticles for Bio-Sensing and Photocatalytic Applications, ACS Omega 2020, 5, 13123−13129

61. D. L. Bish, and J.E. Post, 1989, Modern powder diffraction, Reviews in Mienralogy. Cambridge University Press, Washington, D.C. DOI: https://doi.org/10.1180/claymin.1990.025.4.12

62. D. M. Moore and R. C. Reynolds, Jr. 1997 X-Ray diffraction and the identification and analysis of clay minerals. Oxford University Press, New York, 2nd edn.

63. K. Paulkumar, G. Gnanajobitha, M. Vanaja, S. Rajeshkumar, C. Malarkodi, K. Pandian and G. Annadurai, 2014, *Piper nigrum* leaf and stem assisted green synthesis of silver nanoparticles and evaluation of its antibacterial activity against agricultural plant pathogens. Sci. World J., 2014, 1–9. DOI: http://dx.doi.org/10.1155/2014/829894

64. K. T. Rivero, J. B. Arrieta, N. Fiol, and A. Florido, 2021, Metal and metal oxide nanoparticles: An integrated perspective of the green synthesis methods by natural products and waste valorization: applications and challenges. Comprehensive Analytical Chemistry, ed. S. K. Verma and A. K. Das, Elsevier, 94, ch. 10, 433–469. DOI: https://doi.org/10.1016/bs.coac.2020.12.001.

65. T. W. Bailey, 2013, Antimicrobial assays: comparison of conventional and fluorescence-based methods. J. of Pur. Undergradu. Res., 3. DOI: http://dx.doi.org/10.5703/1288284315146

66. M. Balouiri, M. Sadiki and S. K. Ibnsouda, 2016, Methods for in vitro evaluating antimicrobial activity: A review. J. of Pharm. Anal., 6, 71–79. DOI: http://dx.doi.org/10.1016/j.jpha.2015.11.005

67. S. A. Rupa, M. R. Moni, M. A. M. Patwary, M. M. Mahmud, M. A. Haque, J. Uddin and S. M. T. Abedin, 2022, Synthesis of novel tritopic hydrazone ligands: spectroscopy, biological Activity, DFT, and molecular docking studies. Molecules, 2022, 27, 1–22. DOI: https://doi.org/10.3390/molecules27051656

68. M. M. Rahman, M. B. Islam, M. Biswas and A. H. M. Khurshid Alam, 2015, In vitro antioxidant and free radical scavenging activity of different parts of Tabebuia pallida growing in Bangladesh. BMC Res Notes, 8, 1–9. DOI: https://doi.org/10.1186/s13104-015-1618-6

69. H. Hamrayev and K. Shameli, 2021, Biopolymer-based green synthesis of zinc oxide (ZnO) nanoparticles. IOP Conf. Series: Mat. Sci. and Eng., 1051. DOI: https://doi.org/10.1088/1757-899X/1051/1/012088

70. . K. R. Ahammed, M. Ashaduzzaman, S. C. Paul, M. R. Nath, S. Bhowmik, O. Saha, M. M. Rahaman, S. Bhowmik and T. D. Aka, 2020, Microwave assisted synthesis of zinc oxide (ZnO) nanoparticles in a noble approach: utilization for antibacterial and photocatalytic activity. SN Appl. Sci., 2020, 2. DOI: https://doi.org/10.1007/s42452-020-2762-8

71. H. Çolak and E. Karakose, 2017, Green synthesis and characterization of nanostructured ZnO thin films using *Citrus aurantifolia* (lemon) peel extract by spin-coating method J. Alloys. Compd., 690. DOI: 10.1016/j.jallcom.2016.08.090

72. S.R. Senthilkumar and T. Sivakumar, 2014, Green tea (*Camellia sinensis*) mediated synthesis of Zinc Oxide nanoparticles and studies on their antimicrobial activities. Int. J. Pharm. Pharm. Sci., 2014, 6, 461–465.

73. R. K. Shah, F. Boruah and N. Parween, 2015, Synthesis and characterization of ZnO nanoparticles using leaf extract of *Camellia sinesis* and evaluation of their antimicrobial efficacy Int. J. Curr. Microbiol. Appl. Sci., 2015, 4, 444–450.

74. M. M. N. Yung, C. Mouneyrac, K. M. Y. Leung, 2014, Ecotoxicity of Zinc Oxide Nanoparticles in the Marine Environment. Encyclo. of Nanotech., 1–17. DOI: 10.1007/978-94-007-6178-0_100970-1

75. K. Akhil and S. S. Khan, 2017, Effect of humic acid on the toxicity of bare and capped ZnO nanoparticles on bacteria, algal and crustacean systems. J Photochem Photobio B, 167, 136–149. DOI: http://dx.doi.org/10.1016/j.jphotobiol.2016.12.010

76. K. Davis, R. Yarbrough, M. Froeschle, J. Whitea and H. Rathnayake, 2019, Band gap engineered zinc oxide nanostructures via a sol–gel synthesis of solvent driven shapecontrolled crystal growth. RSC Adv., 9, 14638–14648. DOI: https://doi.org/10.1039/C9RA02091H

77. D. Suresh, P.C. Nethravathi, Udayabhanu, H. Rajanaika, H. Nagabhushana, S.C. Sharma, 2015, Green synthesis of multifunctional zinc oxide (ZnO) nanoparticles using *Cassia fistula* plant extract and their photodegradative, antioxidant and anti-bacterial activities. Mater. Sci. Semicond. Proc., 31, 446–454. DOI: 10.1016/j.mssp.2014.12.023

78. S. S. Rad, A. M. Sani, S. Mohseni, Biosynthesis, characterization and antimicrobial activities of zinc oxide nanoparticles from leaf extract of *Mentha pulegium* (L.), Microbial Pathogenesis 131 (2019) 239–245. https://doi.org/10.1016/j.micpath.2019.04.022

79. E. Selvarajan and V. Mohanasrinivasan, 2013, Biosynthesis and characterization of ZnO nanoparticles using *Lactobacillus plantarum* VITES07. Mater Lett, 112, 180–182. DOI: http://dx.doi.org/10.1016/j.matlet.2013.09.020

80. A. K. M. A. Ullah, M. F. Kabir, M. Akter, A. N. Tamanna, A. Hossain, A. R. M. Tareq, M. N. I. Khan, A. F. Kibria, M. Kurasaki, and M. M. Rahman, 2018, Green synthesis of bio-molecule encapsulated magnetic silver nanoparticles and their antibacterial activity. RSC advances, 8, 37176–37183, DOI: https://doi.org/10.1039/C8RA06908E

81. M. M. Priya, B. K. Selvi, J. A. J. Paul, 2011, Green synthesis of silver nanoparticles from the leaf extract of *Euphobia hirta* and *Nerium indicum*. Dig. J of Nano. and Biostruc., 6, 869–877.

82. J. Y. Song and B. S. Kim, 2008, Biological synthesis of bimetallic Au/Ag nanoparticles using Persimmon (*Diopyros kaki*) leaf extract. Korean. J. chem. Eng., 25, 808–811.

83. S. Jiang, K. Lin and M. Cai, 2020, ZnO nanomaterials: current advancements in antibacterial mechanisms and applications. Front. Chem., 2020, 8, 1–5. doi: 10.3389/fchem.2020.00580

84. A. Roy, O. Bulut, S. Some, A. K. Mandal and M. D. Yilmaz, 2019, Green synthesis of silver nanoparticles: biomolecule-nanoparticle organizations targeting antimicrobial activity. RSC Adv., 2019, 9, 2673–2702, DOI: 10.1039/c8ra08982e

85. J. Guarro, M. I. A. Ayala, J. Gené, J. G. Calzada, C. N. Díez and M. Ortoneda, 1999, Fatal Case of Trichoderma harzianum Infection in a Renal Transplant Recipient. J Clin Microbiol., 37, 3751–3755. doi: 10.1128/jcm.37.11.3751-3755.1999

86. H. Jan, S. A. Shah, S. Shah, A. Khan, M. T. Akbar, M. Rizwan, F. Jan, Wajidullah, N. Akhtar, A. Khattak, and S. Syed, 2021, Green synthesis of Zinc Oxide (ZnO) nanoparticles using aqueous fruit extracts of *Myristica fragrans*: their characterizations and biological and environmental applications. ACS Omega, 6, 9709−9722, DOI: https://doi.org/10.1021/acsomega.1c00310

87. J. Lua, I. Batjikh, J. Hurh, Y. Han, H. Ali, R. Mathiyalaganb, C. Ling, C. J. Ahn, D. C. Yang, 2019, Photocatalytic degradation of methylene blue using biosynthesized Zinc Oxide nanoparticles from bark extract of *Kalopanax septemlobus*. Inter J. Light Electron Optics, 182, 980–985, DOI: https://doi.org/10.1016/j.ijleo.2018.12.016

88 J. Osuntokun, D. C. Onwudiwe and E. E. Ebenso, 2019, Green synthesis of ZnO nanoparticles using aqueous Brassica oleracea L. var. italica and the photocatalytic activity. Green Chemistry Letters and Reviews, 12, 444–457, DOI:10.1080/17518253.2019.1687761

89. M. Murali, N. Kalegowda, H. G. Gowtham, M. A. Ansari, M. N. Alomary, S. Alghamdi, N. Shilpa, S. B. Singh, M. C. Thriveni, M. Aiyaz, N. Angaswamy, N. Lakshmidevi, S. F. Adil, M. R. Hatshan and K. N. Amruthesh, 2021, Plant-Mediated Zinc Oxide nanoparticles: advances in the new millennium towards understanding their therapeutic role in biomedical applications. Pharmaceutics, 13, 1–44, DOI: https://doi.org/10.3390/pharmaceutics13101662

90. H.R. Madan, S.C. Sharma, Udayabhanu, D. Suresh, Y.S. Vidya, H. Nagabhushana, H. Rajanaik, K.S. Anantharaju, S.C. Prashantha and P. S. Maiya, 2015, Facile green fabrication of nanostructure ZnO plates, bullets, flower, prismatic tip, closed pinecone: their antibacterial, antioxidant, photoluminescent and photocatalytic properties. Spectroc Acta Part A: Mol. and Biomol. Spectro. DOI: http://dx.doi.org/10.1016/j.saa.2015.07.067

91. D. Das, B. Chandra, N. P. Phukonc, A. Kalitaa and S. K. Doluia, 2013, Synthesis of ZnO nanoparticles and evaluation of antioxidant and cytotoxic activity. Colloids Surf. B: Biointerfaces, 111, 556–560. DOI: 10.1016/j.colsurfb.2013.06.041

